# *Plasmodium falciparum* schizont stage transcriptome variation among clinical isolates and laboratory-adapted clones

**DOI:** 10.1101/329532

**Authors:** Sarah J Tarr, Ofelia Díaz-Ingelmo, Lindsay B Stewart, Suzanne E Hocking, Lee Murray, Craig W Duffy, Thomas D Otto, Lia Chappell, Julian C Rayner, Gordon A Awandare, David J Conway

## Abstract

Malaria parasite genes exhibit variation in both sequence and expression level. There is much information on sequence polymorphism, but less resolution on natural variation in transcriptomes of parasites at specific developmental stages. This is largely because it is challenging to obtain highly replicated sampling of transcriptomes to overcome potentially confounding technical and biological variation. We address the issue in the major human parasite *Plasmodium falciparum* by obtaining RNA-seq profiles of multiple independent replicate preparations of mature schizont-stage parasites from a panel of clinical isolates recently established in culture and from long-term laboratory-adapted clones. With a goal of robustly identifying variably expressed genes, we show that increasing the numbers of biological sample replicates greatly improves the discovery rate. Generally, six independent replicates of each parasite culture is recommendable as being significantly to lower numbers, although for highly expressed genes variable expression can be detected when fewer replicates are available. A broad comparison identifies genes differing in relative expression between cultured clinical isolates and laboratory-adapted clones. Genes more highly expressed in the laboratory-adapted clones include an AP2 transcription factor gene Pf3D7_0420300 and putative methyl transferase genes. The variable expression of several known merozoite invasion ligands is confirmed, and previously uncharacterised genes are shown to be differentially expressed among clinical isolates. New RT-qPCR assays validate the variation in transcript levels of these genes, and allow quantitation of expression to be extended to a wider panel of clinical isolate samples. These variably expressed genes are new candidates for investigation as potential determinants of alternative parasite developmental pathways or targets of immunity.

**Author summary:** Understanding parasite diversity and adaptation may require characterisation of gene expression variation, and is vital if chemotherapeutic or vaccine development is to consider new candidate targets, but it is technically challenging to generate precise data on clinical isolates. Here, we analyse the transcriptomes of mature *Plasmodium falciparum* schizonts using RNA-sequencing, using large numbers of biological replicate samples to minimise the impact of inter-replicate variation on observed patterns of differential expression. This identifies genes that are differentially expressed in long term laboratory-adapted parasites and recently cultured clinical isolates, as well as among different clinical isolates. In additional samples of schizonts grown in the first cycle *ex vivo* prior to any erythrocyte invasion, expression levels of a selected panel of these genes vary among isolates, but mean levels are similar to those in the continuously cultured clinical isolates, indicating that the latter are useful for experimental studies requiring biological replication.

## Introduction

Variation in gene expression is key to survival and reproduction of microbes, affecting diverse phenotypes such as sexual differentiation [1], adaptation to different growth conditions [2] and evasion of host immunity [3]. Malaria occurs as parasites undergo cycles of invasion and asexual replication inside erythrocytes. Towards the end of each cycle schizonts develop which contain multiple merozoites that need to burst from the host cell and invade a new erythrocyte. The transcriptional program is highly synchronised throughout the asexual parasite replication cycle in erythrocytes [4], but some variation exists between parasite clones [5,6]. Analysis of naturally occurring sequence polymorphism in *Plasmodium falciparum* has shown that selection maintains multiple alleles of many merozoite-stage genes [7,8], some of which encode targets of naturally acquired immunity [9–11]. Genes expressed at this stage can also exhibit pronounced variation in transcription among parasites [7,12–14], although the extent of this variation among laboratory-adapted parasite lines and clinical isolates has not been deeply surveyed.

Accurate quantitation of differential gene expression between organisms or groups of organisms in whole-transcriptome analyses requires biological replicate sampling [15,16]. The importance of minimising sampling error in order to detect real differential gene expression is well recognised [17], and some bioinformatic tools can guide the determination of replicate numbers appropriate to experimental designs [18,19]. In transcriptomic studies of malaria parasites, achieving large numbers of replicate sample preparations is difficult, particularly from clinical isolates [5], and most current understanding is derived from a small number of parasite lines that have been culture-adapted for many years. Examination of genome sequences has identified premature stop codon mutants affecting some transcriptional factor genes and cell signalling protein genes [20,21], as well as particular gene deletions and amplifications only observed in parasites adapted to *in vitro* culture [22,23]. Although culture-adapted parasites are phenotypically and transcriptionally diverse [6,12,24], it is not clear to what extent they reflect the diversity of parasites causing clinical malaria. Parasites may be studied in the first cycle of *ex vivo* growth following isolation from patients, and maturation to schizont stages shows variable expression of invasion-related genes [7,13,14], but a lack of biological replicate measurements limits the precision of whole transcriptome analysis of such samples.

Here we present gene expression profiles of schizont-stage malaria parasites from multiple cultured clinical isolates and laboratory-adapted lines. We conduct RNA-seq analysis with large numbers of replicates per sample, and show that high numbers of biological replicates improves the true-positive discovery rate for identifying differentially expressed genes. This identifies schizont-stage genes that are differentially expressed between laboratory-adapted and cultured clinical isolates, as well as those that are variably expressed among cultured clinical isolates. The results confirm variable transcription in particular genes encoding ligands involved in erythrocyte invasion, and variation in expression of genes that have been less characterised was quantitatively validated by targeted RT-qPCR assays, allowing analysis to be extended to additional isolates from which material is more limited. The expression levels of these genes in a panel of clinical isolates sampled at schizont stage in the first *ex vivo* cycle were similar on average to those observed in cultured clinical isolates. This indicates that cultured parasite lines are useful for studying expression of most genes, although a minority of genes are affected by culture adaptation.

## Results

### Replicate samples of schizont-stage transcriptomes from *P. falciparum* laboratory-adapted clones and clinical isolates

Nineteen biological replicate preparations of schizont-stage parasites were made from cultured *P. falciparum* clone 3D7, and between five and ten biological replicate preparations were made from each of three other laboratory-adapted clones of diverse origin and six clinical isolates from Ghana. Transcriptomes were obtained by sequencing of oligo-dT-primed cDNA (RNA-seq), and mapping reads to the 3D7 reference genome sequence. To minimize potential effects of sequence differences between the different parasites, regions of genes with high levels of allelic polymorphism were masked, so that for polymorphic genes only reads mapping to conserved portions were analysed (S1 file). Genes that have been described as deleted in some parasites were not excluded, as most individual deletions are very rare and exclusion would result in unnecessary loss of data for genes unaffected in the isolates sampled here, but we chose to rather filter out cases where deletion was apparent in analysis. The subtelomeric large gene families *var, rifin* and stevor were excluded as short read sequence mapping is not generally locus-specific for these, so 5,134 genes were analysed in total. Mapped RNA-seq reads for each gene were converted to fragments per kilobase of transcript per million mapped reads (FPKM). Analysis of four paired replicates of schizont preparations from E64-treated and untreated cultures of clone 3D7 showed that E64 treatment had no major effect, with only a single gene showing log_2_ fold difference > 2 (S2 File).

As multiple biological replicates could not have exactly the same distribution of parasite developmental stages, despite being schizont-enriched, to assess the extent of earlier stage transcripts the distributions of FPKMs for each replicate were correlated with FPKM values in reference RNA-seq data for seven cultured time points over a ~48 hour *P. falciparum* cycle [25]. This identified only a small number of replicates that did not have a maximum Spearman’s correlation with parasites at either 40 or 48 hours post-invasion, and these were excluded from further analyses (Fig 1). Within each isolate, pairwise correlations of FPKMs among replicates correlated highly, with average Spearman’s ρ > 0.95 (Fig 2).

**Figure 1.**
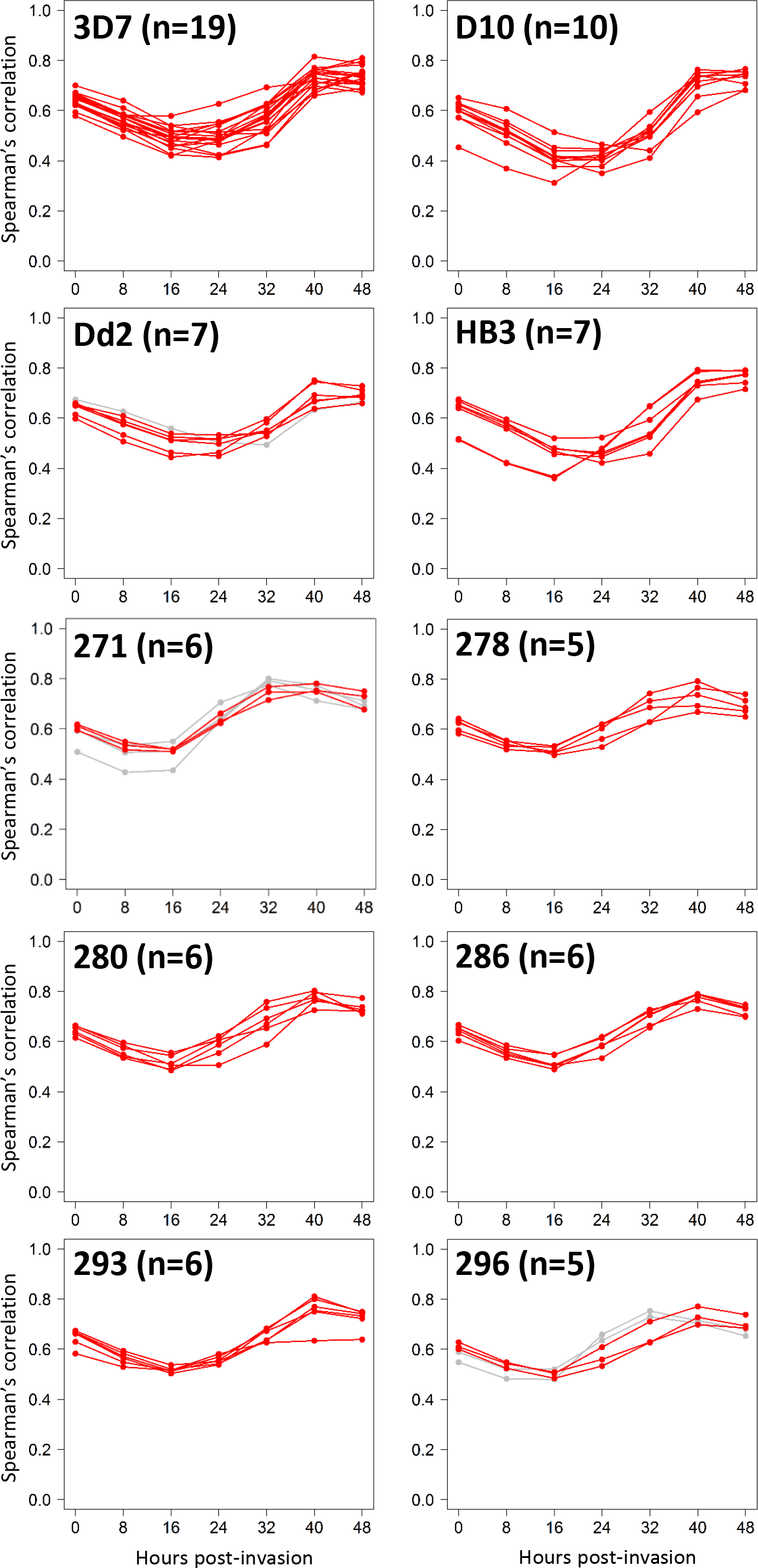
Multiple replicate biological preparations of *P. falciparum* transcriptomes from four long-term laboratory adapted clones and six Ghanaian clinical isolates, showing FPKM values across all genes correlated with previous data from seven time-points across the *P. falciparum* asexual erythrocytic cycle [25]. Each individual parasite culture was enriched for schizont-stage parasites, with egress blocked using E64 treatment for 5.5 hours, and parasites were purified using MACS purification for all of the lines, except for the D10 and HB3 clones and ten of the 3D7 replicates which were purified using Percoll density gradient centrifugation. Red lines plot data for samples with peak correlation at either 40 or 48 hours post-invasion, and grey lines plot replicates that did not maximally correlate with either of these timepoints and which were subsequently excluded from the analysis.

**Figure 2.**
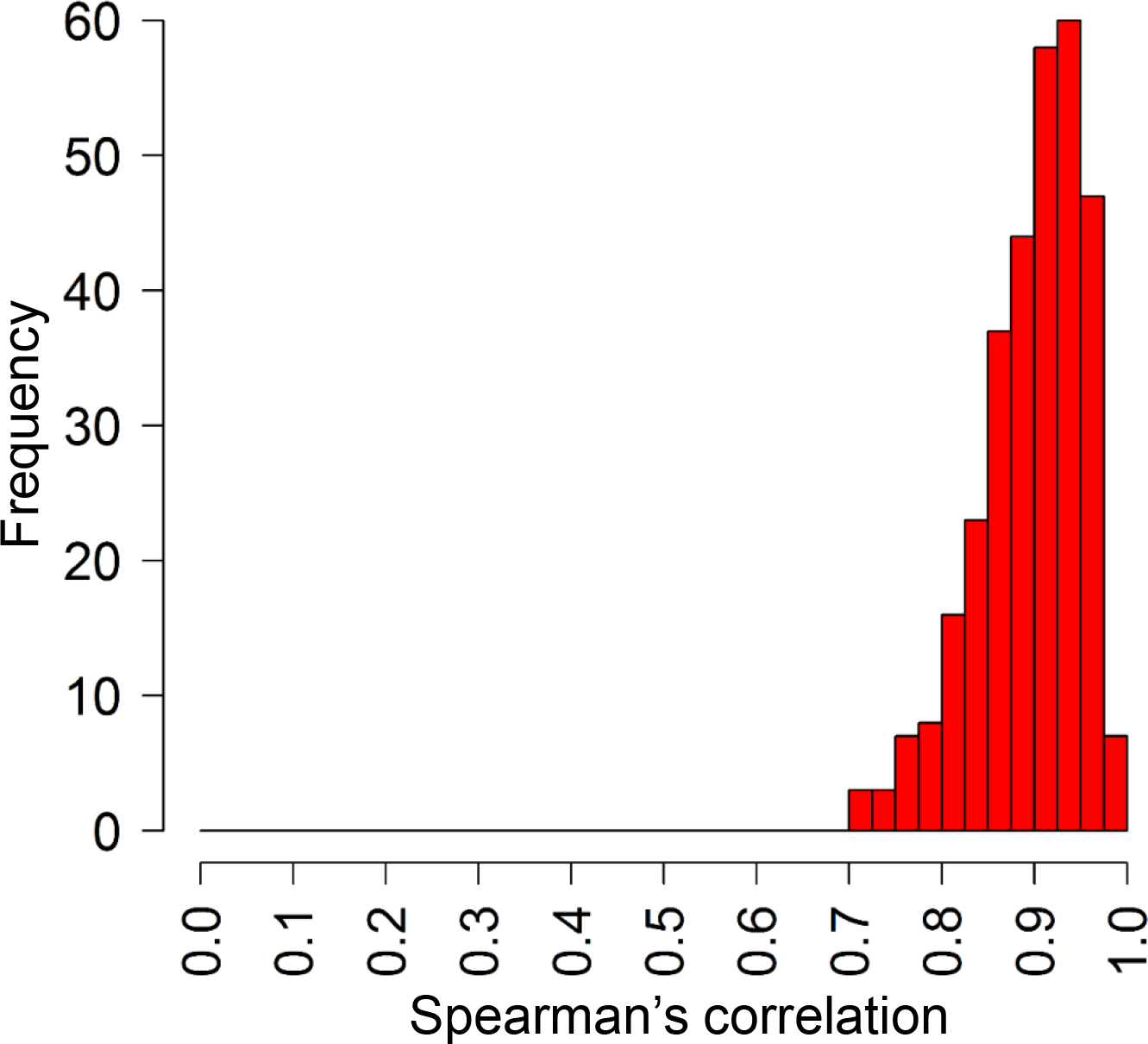
Distribution of correlations of FPKM expression values among biological sample replicate transcriptomes for each isolate or clone. The final numbers of replicates, and median Spearman’s correlation coefficients among replicates were: 3D7 n=19, ρ=0.89; D10 n=10, ρ=0.93; Dd2 n=6, ρ=0.95; HB3 n=7, ρ=0.89; 271 n=3, ρ=0.94; 278 n=5, ρ=0.91; 280 n=6, ρ=0.93; 286 n=6, ρ=0.97; 293 n=6, ρ=0.93; 296 n=3, ρ=0.90.

### Numbers of biological replicates improve detection of differentially expressed genes

We assessed the impact of increasing numbers of biological replicates on detection of genes with differing transcript levels between parasite lines. Data were first compared for the 3D7 and D10 laboratory clones, which each had material from schizonts purified by density gradient centrifugation from ten independent cultures. Comparing all ten replicates of 3D7 and D10 identified 123 genes with log_2_ differences of > 2 in relative transcript levels between the two clones (equivalent to at least 4-fold differences). Achieving this high number of biological replicate samples is difficult, so we assessed what proportion of these 123 genes were captured as being differentially expressed in 100 random samples of two, four, six and eight replicates within each group. As expected, the true-positive proportion of differentially expressed genes detected (using the same statistical criteria of absolute log_2_ fold difference > 2) increased with the number of replicates within each group, median true-positive rates being 0.28, 0.52, 0.67 and 0.79, for two, four, six and eight replicates respectively (Fig 3). While the true-positive rate of genes detected increased with the number of replicates, the median false-positive rates were very low in all cases (among the genes that did not show differences with 10 replicates, the proportions showing apparent differences with two, four, six and eight replicates were 0.001, 0.004, 0.005 and 0.004 respectively).

**Figure 3.**
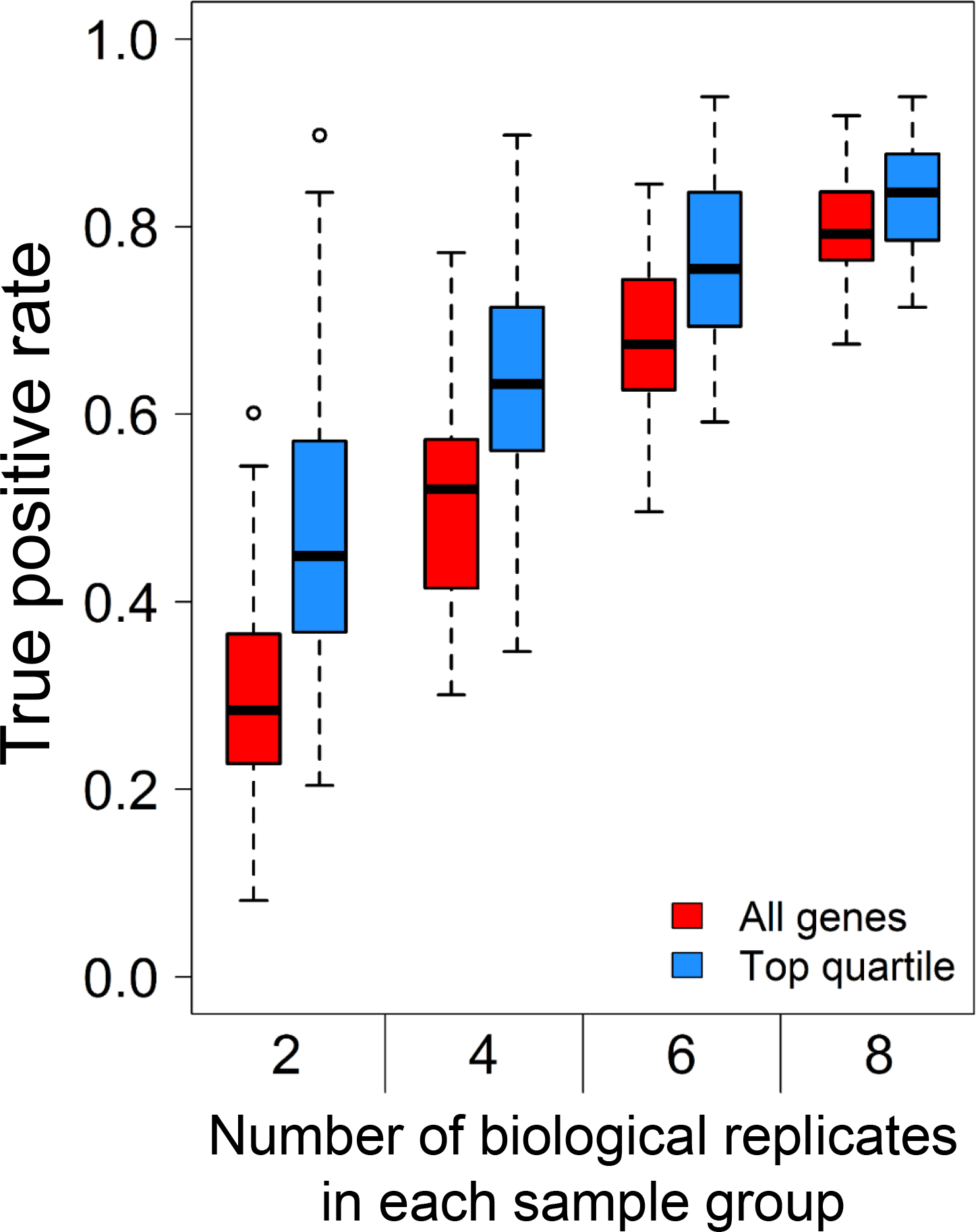
Increased sample replication improves discovery of differentially expressed genes. Assessment of the proportion of genes captured as being differentially expressed between two different parasite clones (3D7 and D10), by taking 100 random samples of two, four, six and eight replicates of each (out of ten initially analysed replicates that identified 123 genes with log_2_ differences of > 2 in relative transcript levels between the two clones).

We also examined the ability to detect differential expression when focusing specifically on genes that are highly expressed in schizont-stage samples, as these are the genes most likely to be functionally important for the merozoite invasive stage. The FPKM values for each sample in each of the 100 comparisons were averaged and the genes with non-zero FPKM values were ranked by their maximum expression level in either strain. Then, the differentially expressed genes that fell within the top quartile of most highly expressed genes in the subsampled data were compared to those identified as differentially expressed among the top quartile of genes in the full analysis. This yielded median true-positive rates of 0.45, 0.63, 0.78 and 0.86 for comparisons respectively containing two, four, six and eight replicates, higher proportions than for the analysis of all genes. Based on these data, it is evident that the use of six independent biological replicates for parasite RNA-seq will detect most genes that are differentially expressed, and that slightly fewer replicates are adequate if analyses only need to focus on highly expressed genes.

### Comparison of gene expression in cultured clinical and laboratory-adapted isolates

Comparing the transcriptomes of long-term laboratory-adapted and cultured clinical isolates, 134 genes (2.6 % of those analysed) had different transcript levels between the two groups (S1 Table). Among the genes within the top quartile of expression values overall, 20 genes (1.6 %) were differentially expressed between the laboratory-adapted and clinical isolate groups, a lower proportion than among the rest of the genes (Odds Ratio 0.52, Fisher’s exact P = 0.006). It is possible that there is a higher background of false positive differences for the less expressed genes, due to lower read coverage giving greater noise in the relative ratios. Examining the 20 genes with differing transcript levels among the top quartile of expressed genes, several were seen to contain a deletion in one of the parasite lines or to be members of multigene families. For example, PF3D7_1371600 (*ebl-1*) and PF3D7_0935700 are deleted in the HB3 strain and D10 strain respectively (Fig 4), as previously documented [26,27]. After excluding these genes and others for which one or more of the samples were presumed to contain genetic deletions, and excluding members of sub-telomeric multi-gene families, ten single-locus genes were identified as showing differential expression between laboratory and clinical isolates (Table 1).

**Figure 4.**
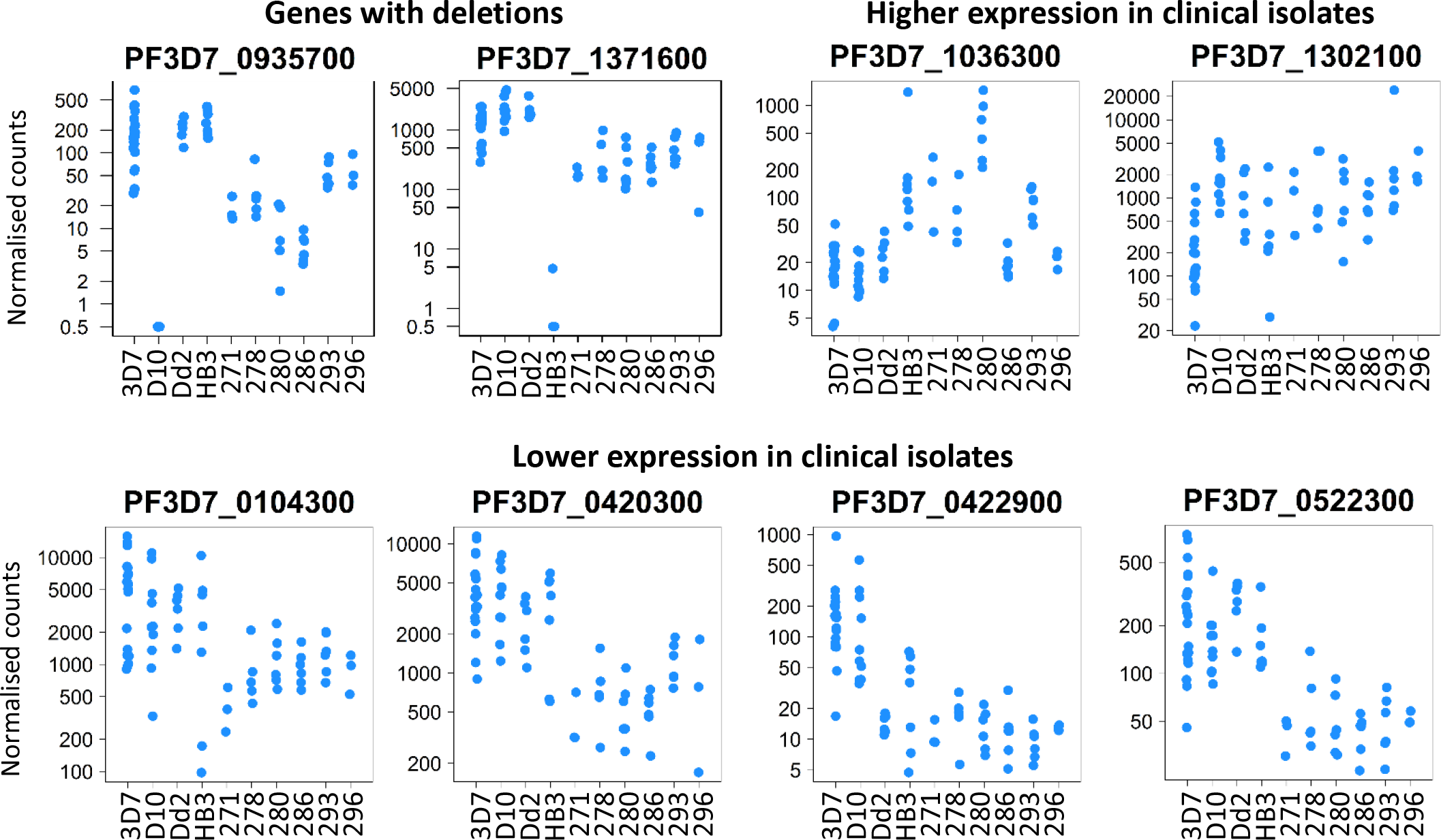
Differential gene transcript levels among preparations of schizont-enriched P. falciparum cultures of laboratory-adapted clones and clinical isolates. Plots of normalised RNA-seq read counts for eight of the genes showing differences between four laboratory-adapted clones and six clinical isolates. Each point shows the value for an independent biological replicate sample. The two top left panels show examples of genes for which one of the laboratory-adapted clones had a deletion (HB3 and D10 for the respective genes). These, and other genes with suspected or known deletions in the sampled parasites, are excluded from the list of differentially expressed genes in Table 1.

Eight of the ten highly expressed genes showing a difference between the groups had higher transcript levels in the long-term laboratory-adapted clones than in the cultured clinical isolates (Table 1). These include an AP2 transcription factor gene PF3D7_0420300, as well as two predicted methyltransferase genes (PF3D7_0422900 and PF3D7_0522300), and gene PF3D7_0104300 encoding ubiquitin-binding protein 1 which is involved in protein turnover [28] (Fig 4). The only two genes that had higher transcript levels in cultured clinical isolates were PF3D7_1036300 which encodes the merozoite surface protein MSPDBL2, and PF3D7_1302100 which encodes the gamete antigen 27/25 (Fig 4). Previous data has shown MSPDBL2 to be variably expressed among *P. falciparum* isolates [7], and experimental analysis has shown transcript levels to be increased in parasite cultures when the heterochromatin protein 1 is suppressed by gametocyte development protein 1 (GDV1) leading to gametocytogenesis [29,30], whereas gamete antigen 27/25 is a known marker of early gametocytogenesis [30,31]. Gametocyte gene transcripts were not generally more highly expressed in the clinical isolates, and gene PF3D7_1327300 that has been previously described to be transcribed in gametocytes [32] showed lower expression.

**Table 1.** *P. falciparum* genes in the top quartile of expression showing a difference in schizont stage transcript levels between long-term laboratory-adapted clones and recent clinical isolates in culture

| Gene ID | Product annotation | Log<sub>2</sub> fold difference | Wald test |
| --- | --- | --- | --- |
| ***Higher transcription in long-term laboratory-adapted clones*** |  |  |  |
| PF3D7_0422900 | methyltransferase, putative | -2.50 | -6.48 |
| PF3D7_0501300 | skeleton-binding protein 1 | -2.50 | -4.98 |
| PF3D7_0410000 | erythrocyte vesicle protein 1 | -2.38 | -10.62 |
| PF3D7_0104300 | ubiquitin carboxyl-terminal hydrolase 1, putative | -2.18 | -6.13 |
| PF3D7_0522300 | 18S rRNA (guanine-N(7))-methyltransferase, putative | -2.15 | -8.67 |
| PF3D7_0420300 | AP2 domain transcription factor, putative | -2.02 | -7.19 |
| PF3D7_1030200 | claudin-like apicomplexan microneme protein, putative | -2.00 | -9.21 |
| PF3D7_1327300 | conserved *Plasmodium* protein, unknown function | -2.00 | -4.97 |
| ***Higher transcription in clinical isolates*** |  |  |  |
| PF3D7_1036300 | duffy binding-like merozoite surface protein 2 | 3.02 | 5.32 |
| PF3D7_1302100 | gamete antigen 27/25 | 2.02 | 3.99 |

We investigated whether there was an overall association between the genes differentially expressed and targets of HP1 regulation [29,33]. Of the HP1-regulated genes previously identified [29], 118 genes are among those analysed here (*var, rifin* and *stevor* gene families were excluded). Of the 134 genes differentially expressed between the two groups, 24 (17.9 %) were targets of HP1 regulation, a higher proportion than among the rest of the genes (Odds Ratio 11.3, Fisher’s exact P = 1.3 × 10^−5^). Among 20 differentially expressed genes in the top quartile of expression values, two (10 %) were targets of HP1 regulation, a higher proportion than among the remaining genes in the top quartile (Odds Ratio 7.6, Fisher’s exact P = 0.037).

### Differential expression of schizont-stage genes among cultured clinical isolates

Overall, 214 genes (4.2 %) showed a log_2_ fold difference > 2 for at least one of the fifteen pairwise comparisons among cultured clinical isolates. Of those genes in the top quartile of expression levels, 39 genes (3.0 %) showed a difference for at least one pairwise comparison, a lower proportion than among the less expressed genes (Odds Ratio 0.66, Fisher’s Exact P = 0.02) (S2 Table). Some of the variably expressed genes encode merozoite proteins previously characterised as transcriptionally variable among *ex vivo* clinical isolates [14], including *dblmsp2* and *msp6*, as well as erythrocyte binding antigen genes *eba-175*, *eba-181*, and *eba-140* (which is also deleted in one of the laboratory adapted clones) (S1 Fig).

There was an over-representation of GDV1-regulated genes [30] among the 214 genes differentially expressed in cultured clinical isolate comparisons (Fisher’s exact test odds ratio 12.5, Fisher’s Exact P = 1.9 × 10^−9^). Of the genes differentially expressed among cultured clinical isolates, 64 % were also differentially expressed in comparisons among the long-term laboratory-adapted clones (Odds Ratio 17.4, Fisher’s exact P = 2.2 × 10^−16^). There was also strong overlap when only considering genes in the top quartile of expression overall, 44 % of genes differentially expressed among cultured clinical isolates also being differentially expressed among laboratory-adapted lines (Odds Ratio 7.9, Fisher’s exact P = 2.4 × 10^−8^).

To validate the data obtained through RNA-seq, a subset of genes was selected for quantitation by reverse-transcription quantitative PCR (RT-qPCR). We chose differentially expressed genes encoding proteins containing predicted transmembrane domains or signal peptides, thereby likely to enter the parasite secretory pathway, excluding genes previously characterised as variably expressed or members of multi-gene families (S2 Table). Eight genes were selected for assay (S2 Fig), two of which encode known antigens (merozoite-associated tryptophan-rich antigen [34] liver-stage antigen-3 [35]). These gene transcripts were quantified by RT-qPCR in 58 RNA preparations from the laboratory clones and clinical isolates that had been analysed by RNA-seq. Transcript levels were normalised to those of a house-keeping gene (serine-tRNA ligase PF3D7_0717700) and correlated with similarly normalised FPKM expression values for the RNA-seq data. Each gene showed strong positive and highly significant correlation between RNA-seq and RT-qPCR-derived expression measures, all except one having correlation coefficients > 0.8 (Fig 5).

**Figure 5.**
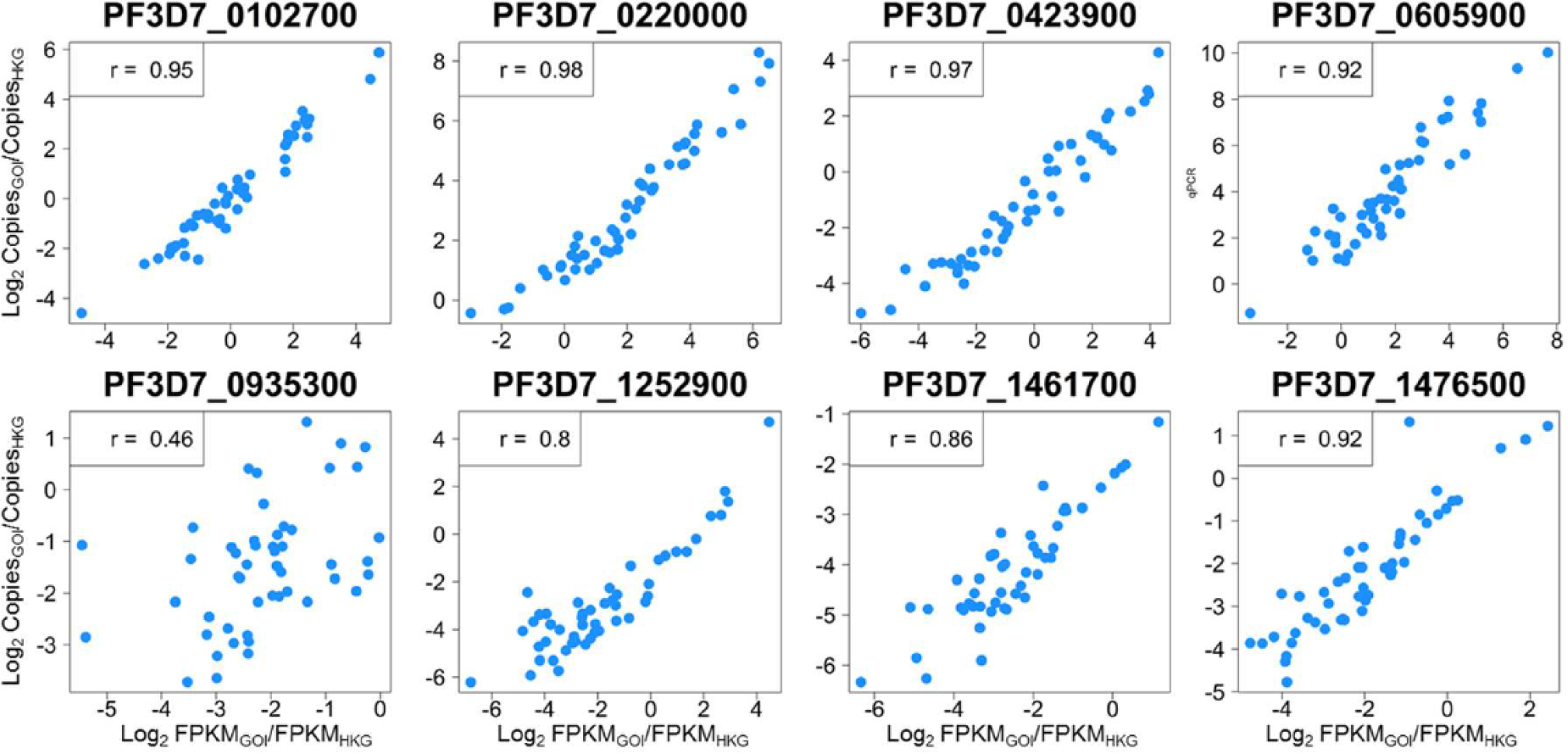
High correlations between RNA-seq and RT-qPCR expression measures. Eight genes identified as differentially expressed among clinical isolates by RNA-seq were validated by RT-qPCR for 49 of the independent RNA preparations from the four laboratory-adapted clones and six cultured clinical isolates under study. Scatter plots for each gene show the gene of interest (GOI) copies (log_2_-transformed) normalised to house-keeping gene copies (HKG; PF3D7_0717700), correlated with GOI FPKM values (log_2_-transformed) normalised to HKG FPKM values. Spearman’s correlation coefficients are shown for each plot. For PF3D7_1461700, one sample had an FPKM value of 0 which was not plotted but is included in the calculation of correlation.

### Variable expression in first *ex vivo* cycle schizonts

We next determined the expression levels in additional clinical samples from Ghana matured during the first cycle *ex vivo* until a large proportion of parasites were schizonts. For nine of these isolates there was sufficient material for RNA-seq to be performed, and results were analysed for seven of these (two isolates were excluded from analysis as whole-transcriptome staging of FPKM values showed maximal correlation with stages earlier than schizonts in a reference time-course). For six isolates grown *ex vivo* to schizont stage there was sufficient material for quantification of the selected eight gene panel by RT-qPCR. Characterisation of the *ex vivo* samples using either method showed considerable variation in relative expression levels of each gene, but the mean levels of each gene across the isolates were similar to those determined in the cultured parasites that had extensive sample replication (Fig 6). Without sample replication of the first cycle *ex vivo* clinical samples, the contribution of random sampling variation and uncontrolled effects on the expression levels cannot be accounted for, but the observation that for each gene the normalised expression values have a similar mean to that for cultured clinical lines indicates no systematic differences in expression between the initial *ex vivo* cycle and under continuous culture conditions. These results support the use of cultured clinical isolates as a means of studying parasite gene expression to reflect that occurring in original populations, while enabling the necessary experimental replication for many analyses.

**Figure 6.**
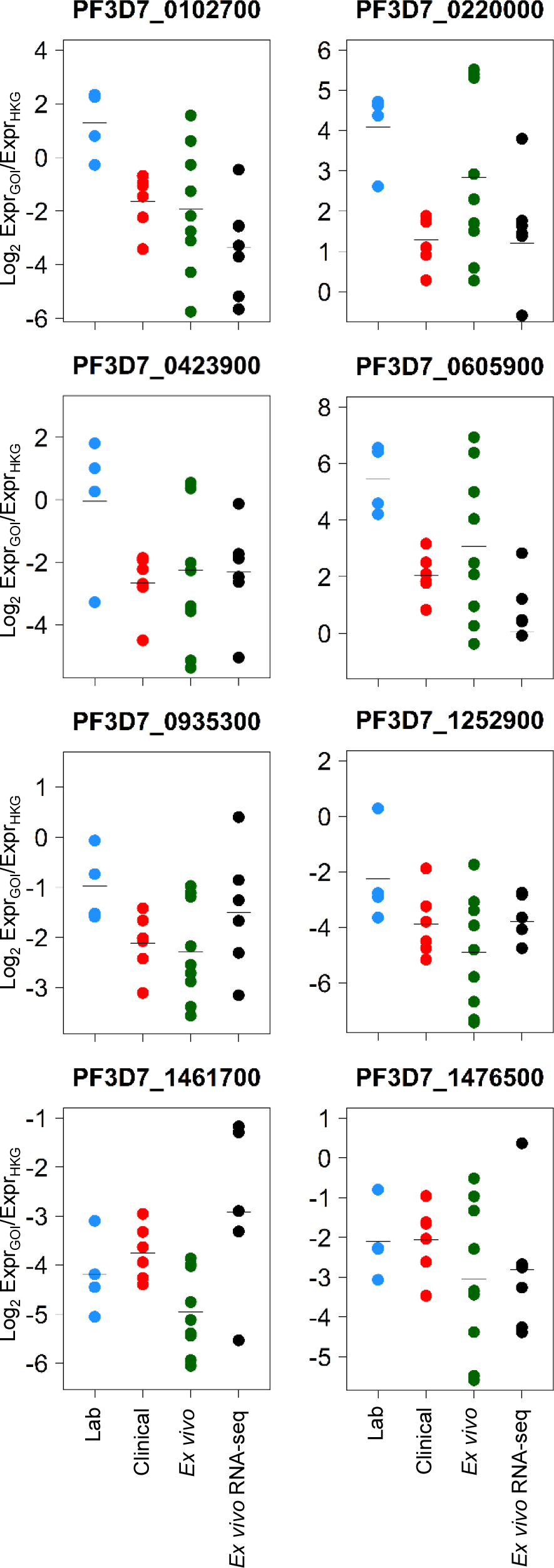
Transcript levels of variably expressed genes for first cycle *ex vivo* clinical samples compared with continuously cultured isolates. For the laboratory and cultured clinical isolates, each point represents the mean log_2_ normalised expression value for multiple replicates of each sample under study. For the ex vivo clinical isolates, each point represents log_2_ normalised expression values for the individual, unreplicated samples under study. Horizontal bars indicate the mean log_2_ normalised expression value for all samples within the group.

## Discussion

Transcriptomic analyses perform optimally with increased levels of sample replication to minimise the impact of stochastic or technical variation in individual samples [17,36]. This is particularly important for studies that focus on a defined developmental stage, in which error can be generated due to subtle differences between samples. However, the nature of clinical samples of malaria parasites is such that replication is technically very difficult. In order to assess the degree of replication useful to robustly identify differentially expressed genes, we first generated RNA-seq profiles from multiple replicates of laboratory-adapted clones, and used the data to calculate the true- and false-discovery rates obtained using fewer replicates. Consistent with studies on other cultured eukaryotes [17], we show that for malaria parasites increasing the sample replication improves the true-positive discovery rate in identifying variably expressed genes, and indicate that six independent replicates are useful to balance accuracy and experimental feasibility. Where these numbers of replicates are not obtainable, it is recommended to focus on genes that are highly expressed in order to achieve as much accuracy as possible per gene.

It has been proposed that spontaneous transcriptional variation within parasite populations is a strategy that ensures fitness of parasites facing a range of potential and changing selective pressures [6], and it is likely that the rates and mechanisms of regulation differ among most genes. In comparison between long-term laboratory-adapted clones and cultured clinical isolates, we found that among the genes most differentially expressed were an AP2-transcription factor [37] and two methyltransferases, the functional roles of which remain to be investigated. Most of the differentially expressed genes had lower transcription in the clinical isolates, suggesting that these may be normally repressed. HP1 is responsible for extensive gene silencing in malaria parasites [30], and we showed an over representation of HP1-regulated genes among those differentially expressed between laboratory-adapted clones and cultured clinical isolates. It would be interesting to undertake further studies directly on HP1 and other chromatin modifications in recent clinical isolates, as experimental studies have so far focused on long-term laboratory-adapted clones. Interestingly, among the top quartile of genes expressed, the only two genes that were more highly expressed in clinical isolates were an early marker of gametocytogenesis (*Pfs25/27*) and a merozoite surface protein gene (*mspdbl2*) encoding a product expressed all merozoites within only a minority of schizonts [7] which may be a marker of parasites committed to sexual development [29]. Studies of gametocytogenesis in clinical isolates will be important to understand the process of commitment to sexual differentiation and transmission, as also indicated by transcriptome comparisons that sampled different stages without having multiple replicates for each stage [5].

Through pairwise comparisons of individual cultured clinical isolates, we identified many genes that were differentially expressed among clinical isolates. Some of the genes identified have previously been shown to be differentially expressed through targeted qRT-PCR assays of schizont-stage *ex vivo* cultured clinical isolates [7,13,14]. Focusing on genes for which variable expression has not been previously studied, we selected a panel of eight highly differentially expressed genes that encode secreted proteins. Of these, the merozoite-associated tryptophan-rich antigen and liver stage antigen 3 have been previously identified as merozoite proteins [34,35], and another gene (PF3D7_0423900) is adjacent to loci encoding the cysteine-rich protective antigen (CyRPA) and reticulocyte binding homologue 5 (Rh5) which are targets of antibodies that inhibit merozoite invasion. We extensively validated the RNA-seq data by RT-qPCR for these genes, and extended quantitation to additional *ex vivo* clinical isolate samples. Despite the lack of sample replicates for the latter, results showed that average expression levels for each gene were similar to those for the cultured samples that had biological replicates.

Together, these data highlight new candidates for investigation as potential determinants of alternative developmental pathways or targets of immunity. Characterisation of gene transcript levels is only one level of analysis, as proteomic characterisation [38] and analysis of phosphorylation and other protein modifications [39,40] may also identify functional determinants, and large-scale analysis of parasite protein variation has not yet been attempted on clinical isolates. A more complete description of transcriptome variation may also be derived in future by analysis of individual infected cells, which has not been undertaken on parasites from clinical isolates, as initial single-cell sequencing studies of laboratory model malaria parasites have only recently begun to probe the expression pathways of different life-cycle stages [41–43]. Scaling up transcriptomic analyses to clinical studies is challenging, and biological replicate preparations of parasite samples may only be feasible in a limited number of cases. In an epidemiological context, reduction of error also requires large sample sizes and tight definition of clinical phenotypes, and more statistical analysis and modelling of covariates may be appropriate, ideally including data on host transcriptome variation within the same samples [44].

## Methods

### Ethical approval

The study of clinical parasite transcriptomes was approved by the Ethics committees of the Ghana Health Service, the Noguchi Memorial Institute for Medical Research, University of Ghana, the Kintampo Health Research Centre, the Navrongo Health Research Centre and the London School of Hygiene and Tropical Medicine. Written informed consent was obtained from parents or legal guardians of all participating children, and additional assent was received from children over 10 years old.

### *P. falciparum* isolates sampled from clinical malaria cases

Blood samples were collected from clinical malaria cases attending Ghana government health facilities between 2012 and 2013, in Kintampo (Brong-Ahafo Region of central Ghana), and Navrongo (Kassena-Nankana East Municipality, in the Upper East Region of northern Ghana). Patients were eligible to participate in the study if they had uncomplicated clinical malaria, were aged 2-14 years, tested positive for *P. falciparum* malaria by lateral flow rapid diagnostic test and slide microscopy, and had not taken antimalarial drugs during the 72 hours preceding sample collection. Venous blood samples (up to 5 ml) were collected into heparinized Vacutainer tubes (BD Biosciences). Blood samples were centrifuged, plasma separated and leukocyte buffy coat cell layer removed, and erythrocytes were cryopreserved in glycerolyte and stored frozen at −80°C or in liquid nitrogen until shipment on dry ice to the London School of Hygiene and Tropical Medicine.

### Parasite culture and enrichment for schizont stages

Parasites from *ex vivo* clinical samples were matured within the original patient erythrocytes at 2-5 % haematoctit for up to 48 h in RPMI 1640 medium containing 2 % human AB serum (GE Healthcare) and 0.3 % Albumax II (Thermo Fisher Scientific) under an atmosphere of 5 % O_2_, 5 % CO_2_, and 90 % N_2_ at 37 °C, until most parasites were at the schizont stage of parasite development, at which point RNA was extracted directly from the bulk cultures. Laboratory-adapted clones and cultured clinical isolates were grown in erythrocytes at 2-5 % haematocrit in RPMI 1640 medium containing 0.5 % Albumax II, at 37 °C. Laboratory-adapted clones were maintained under atmospheric air with 5 % CO_2_, and cultured clinical isolates were maintained in 5 % O_2_, 5 % CO_2_, 90 % N_2_. Schizonts were isolated by either MACS or discontinuous density centrifugation methods.

For MACS purification of cultured clinical isolates, parasites were prepared from 100 ml cultures with at least 0.7 % of erythrocytes containing schizonts. Parasites were isolated by magnetic purification using magnetic LD Separation columns (Miltenyi Biotech). One column was used per 25 ml culture, and washed twice in 3 ml of culture medium at room temperature. Parasite culture was pelleted at 500 g for 5 minutes, then re-suspended in 3 ml culture medium per 1 ml of packed cell volume. The re-suspended material was bound to the MACS column, which was then washed three times with 3 ml culture medium, and schizonts were eluted twice by removing the magnet from the column and forcing 2 ml culture medium through the column into a 15 ml collection tube. Finally, the schizonts were pelleted by centrifugation at 500 g for 5 minutes and the pellet volume was estimated, with 0.5 μl used for Giemsa-stained microscopical examination to assess staging, 1 μl added back to 250 μl culture at 0.8 % hematocrit to follow the progression. Remaining parasites were re-suspended in 1.5 ml of culture medium with 10 μM E64 in a 12 well plate, and parasites were incubated for 5.5 hours in 5 % CO_2_ at 37 °C, before centrifugation in a 1.5 ml tube and processing for RNA extraction.

For discontinuous density centrifugation purification, parasites were maintained as 25 ml cultures at 2.5 % hematocrit. Cultures were used when > 1 % erythrocytes contained parasites with multiple nuclei. Schizonts were purified using 70 % Percoll (GE Healthcare)/2.93 % sorbitol/PBS overlayed with 35 % Percoll/1.47 % sorbitol/PBS, which was overlayed with parasitized erythrocytes, allowing the schizonts to be separated by centrifugation at 2500 g for 10 minutes at 24 °C, with a light brake at the end of centrifugation. Purified schizonts were washed once in complete medium and the pellet volume was estimated. Six pellet volumes of 50 % haematocrit erythrocytes were added to the pellet and resuspended for slide examination. The sample was smeared, and resuspended in 6 ml of complete culture medium. Of this, 1 ml was used as a control untreated sample to track parasite egress. E64 was added to the remaining 5 ml at a final conentraction of 10 μM. Control and E64-treated samples were placed at 37 °C, 5 % CO_2_ static incubator for 5.5 hours. Schizonts from the E64-treated culture were overlaid on 70 % Percoll/2.93 % sorbitol/PBS and separated by centrifugation at 2500 g for 10 minutes at 24 °C, with a light brake at the end of centrifugation. The schizont layer was washed once in complete culture medium and final cell pellets of approximately 10 - 20 μl were used for RNA extraction.

### RNA extraction

Pellets were resuspended 500 μl in TRIzol^®^ reagent (Thermo Fisher Scientific) and stored at −80 °C until RNA extraction following manufacturer’s instructions. Final RNA pellets were resuspended in 100 μl RNase-free H_2_O. A second RNA clean-up and on-column DNase treatment was carried out using RNeasy mini columns (Qiagen), with RNA eluted in 30-50 μl RNase-free H_2_O and concentration quantified by Qubit High Sensitivity RNA Assay (Thermo Fisher Scientific). Samples containing at least 500 ng RNA were considered for analysis after the RNA integrity was checked on an Agilent Bioanalyzer using RNA 6000 Nano reagents and chips (Agilent Genomics).

### RNA-seq library preparation and sequencing

Laboratory isolate and cultured clinical isolate RNA-seq libraries were prepared using TruSeq Stranded mRNA Library Prep Kit (Illumina) using 500 ng - 1 μg RNA as per the Illumina TruSeq Stranded mRNA protocol for MiSeq sequencers. Libraries were validated on an Agilent Bioanalyzer using DNA 1000 reagents and chips (Agilent Genomics) to quantify library sizes and confirm the absence of primer dimers. Libraries were quantified using a KAPA Universal Library Quantification kit (Roche Diagnostics Limited) on a 7500 Fast Real-Time PCR System (Thermo Fisher Scientific) and library concentrations were adjusted for library size. 12 - 15 pM pooled libraries were sequenced on a MiSeq System (Illumina) using a MiSeq Reagent Kit v3 (Illumina) with 2 × 75 cycles.

*Ex vivo* schizont-enriched *P. falciparum* isolate RNA-seq libraries were prepared using a modified protocol. PolyA+ RNA (mRNA) was selected using magnetic oligo-d(T) beads. mRNA was reverse transcribed using Superscript III^®^ (Thermo Fisher Scientific), primed using oligo-d(T) primers. dUTP was included during second-strand cDNA synthesis. The resulting double stranded cDNA was fragmented using a Covaris AFA sonicator. Sheared double stranded cDNA was dA-tailed, end repaired, and “PCR-free” barcoded sequencing adaptors (Bioo Scientific) [45] were ligated (NEB). Libraries were cleaned up twice, using solid phase reversible immobilisation beads, and eluted in EB buffer (Qiagen). Second strand cDNA was removed using uracil-specific excision reagent enzyme mix (NEB). Libraries were quantified by quantitative PCR prior to sequencing on an Illumina MiSeq sequencer.

### Data curation and analysis

Raw Illumina sequence reads were aligned to the *P. falciparum* 3D7 v3 genome and converted to ‘bam’ format using samtools [46]. Reads with MAPQ scores < 60 were removed. Reads were counted using the “summarizeOverlaps” feature of the GenomicAlignments package [47] in R, against a previously published *P. falciparum* genome annotation file that had been masked for regions of polymorphism (detailed in S2 File) to remove known regions of genomic polymorphism, highly polymorphic gene families and duplicated genes.

Fragments Per Kilobase of transcript per Million mapped reads (FPKMs) for the data here and those of previously published data on a developmental time course of the clone 3D7 [25] were calculated using the ‘fpkm’ function of DESeq2 [48] in R. FPKMs for all genes in each of our parasite preparations were correlated using a Spearman’s Rank correlation with FPKMs for each of the seven time points (0, 8, 16, 24, 32, 40 and 48 hours post-invasion; S1 Table). Replicates with peak Spearman correlation values of ρ > 0.7 at the latest time points (40 or 48 hours) were included for further analysis. Differential expression analysis was conducted using DESeq2 in R.

### RT-PCR assays

150 - 500 ng total RNA from each preparation of parasite schizonts was reverse transcribed using Superscript II^®^ (Thermo Fisher Scientific) with 250 ng random hexamer oligonucleotide primers per 20 μl reaction. Quantitative PCR (qPCR) was carried out using SYBR^®^ Select Master Mix (Thermo Fisher Scientific) with 500 nM forward and reverse primers, in a Prism 7500 Fast qPCR machine. For each gene, threshold-cycle values were quantified against a serially diluted genomic DNA (Dd2 strain) standard curve, run on the same plate. Cycling parameters were: 50°C for 2 minutes, 95°C for 2 minutes followed by 40 cycles of 95 °C for 15 seconds and 60 °C for 1 minute. All wells were run as 10 μl volumes in technical duplicate. The qPCR copy numbers were normalised against copies of a house-keeping gene, PF3D7_0717700 [49]. PCR primer pair sequences are as follows: PF3D7_0102700 5’-CAACCAGACAAACAACAAGAAA-3’ and 5’-TCCATTCTGATGCTTTCCAC-3’, PF3D7_0220000 5’-GTAAATGGTGAATTAGCTAGTGAAG-3’ and 5’- CCTTTATATCTTCGGCTTCTTCT-3’, PF3D7_0423900 5’-GAGAAGAAGCCAAAGTAAATGAAC-3’ and 5’-GAATGTGTCAAGTGCATCATAA-3’, PF3D7_0605900 5’-CGCAATAACAAGAAGTCACAAA-3’ and 5’-AAGATTGTAGGAGGGTGTTAAT-3’, PF3D7_0935300 5’-GGGCTGCTGTTATACCTTG-3’ and 5’-AGAATGAGGAGGTTCAGTTTG-3’, PF3D7_1252900 5’-CCTTAGTAGAACTTCAAAGTGACA-3’ and 5’-TGTAACCAACTACGAAATTCCC-3’, PF3D7_1461700 5’-TGCTGACTTTCAAGAGATAAGG-3’ and 5’-TTTCATTTGTTCATTTTGTACAACC-3’, PF3D7_1476500 5’-CTTCGATTCACAAATGCAGAAA-3’ and 5’-CGTATTCCACACCTTGTTCTAT-3’, PF3D7_0717700 5’-AAGTAGCAGGTCATCGTGGTT-3’ and 5’-GTTCGGCACATTCTTCCATAA-3’.

### Data open access

All *P. falciparum* RNA-seq data from the laboratory-adapted and cultured clinical isolates have been submitted for open access at the Gene Expression Omnibus (https://www.ncbi.nlm.nih.gov/geo/), entry: GSE113718. The *P. falciparum* RNA-seq data from the first cycle *ex vivo* isolates have been submitted to the European Nucleotide Accession Archive (https://www.ebi.ac.uk/ena), study: ERP103955.

## Acknowledgements

We are grateful to malaria patients who contributed blood samples for analysis of parasite transcription, and to colleagues who supported the clinical sample collection and initial laboratory processing. We thank Alistair Miles and Antoine Claessens for help with identification of genomic coordinates containing highly polymorphic sequences within genes.

## Supplementary Material

**S1 Figure. Differential expression of merozoite invasion-related genes among schizonts from different parasite cultures.** Distributions of read counts (normalised to library size) for eight genes, for replicated laboratory-adapted and clinical isolate samples, showing data from each replicate culture preparation of each strain.

**S2 Figure. Gene expression levels for eight genes differentially expressed among clinical isolates.** Distributions of read counts (normalised to library size) for eight genes, showing data from each replicate culture preparation of each strain.

**S1 Table.** Genes differentially expressed between clinical and laboratory lines by absolute log2 fold > 2 (at least 4 fold difference on average). For genes among the top quartile of expression values genome-wide (top 18 genes in the table), members of multigene families and genes in which strain-specific deletions may be responsible for differences are annotated with asterisks (*).

**S2 Table.** Log_2_ fold changes of genes differentially expressed among pairwise comparisons of six cultured clinical isolates with multiple schizont preparations of each, among genes within the top quartile of expression overall.

**S1 File.** Masking of *P. falciparum* genome annotation file for polymorphic regions of genes.

**S2 File.** Analysis of paired E64-treated and untreated *P. falciparum* 3D7 schizonts

**S3 File.** Sequence annotation file Pf3D7.May2015.NoSplice.LSHTM.gtf (see S1 Table for details)

**S4 File.** Sequence annotation file GTF_VarRifStev_filtered out.gtf (see S1 Table for details)

